# Synthesis, anticancer properties, and biological profiling of synthetic glycan analogs of proscillaridin A

**DOI:** 10.1101/2025.03.31.646140

**Authors:** Shreya Somani, Lekhya Menta, Feng Wang-Johanning, Gary Johanning, Seoyeon Hong, Edward Njoo

## Abstract

Several cardiac glycosides, including digoxin, digitoxin, and proscillaridin A, have been originally identified as cardiomyocyte modulators and are currently being investigated for their anticancer properties. These cardiac glycosides are generally classified into cardenolides and bufadienolides, which bear butenolide and pyrone D-ring functionality, respectively, and have exhibited remarkable *in vitro* toxicity in various cancerous cell lines. As simple modifications on steroidal small molecules have demonstrated success in augmenting bioavailability or enhancing downstream biological activities, we sought to prepare synthetic analogs of proscillaridin A, a bufadienolide isolated from the genus *Scilla*. We synthesized two novel analogs of proscillaridin A bearing acetate esters or a dimethyl ketal to investigate how strategies of ketalization or acetylation of the A-ring allylic glycoside might alter its anticancer properties. The antiproliferative activity of these compounds was evaluated alongside proscillaridin A and two model cardiac glycosides, digoxin and digitoxin, across several colorectal and liver cancer cell lines. Through a diverse panel of cell viability and cytotoxicity experiments, reporter assays, and cell cycle and protein marker analysis by flow cytometry, we find that ketalization of the glycan of proscillaridin A provides similar, and in some cases enhanced, *in vitro* potency. This study establishes the foundation for current and further *in vitro* and *in vivo* evaluation of glycan analogs of proscillaridin A.

## Introduction

Cardiac glycosides are a class of glycosylated steroidal natural products isolated from phytochemical sources including plants of the genera *Digitalis* and *Scilla* and have been investigated for their potent biological activity and unique therapeutic potential [1–3]. These compounds, which putatively function as antifeedants in their source species [4–6], share a steroidal tetracarbocyclic core but differ in the D-ring lactone and A-ring glycan **(Fig. 1)**. Cardenolides, such as digoxin **1** from the *Digitalis* genus, contain a 5-membered butenolide lactone ring, and bufadienolides, such as proscillaridin A **3** from the *Scilla* genus, contain a D-ring 6-membered pyrone ring [7].

**Fig. 1.**
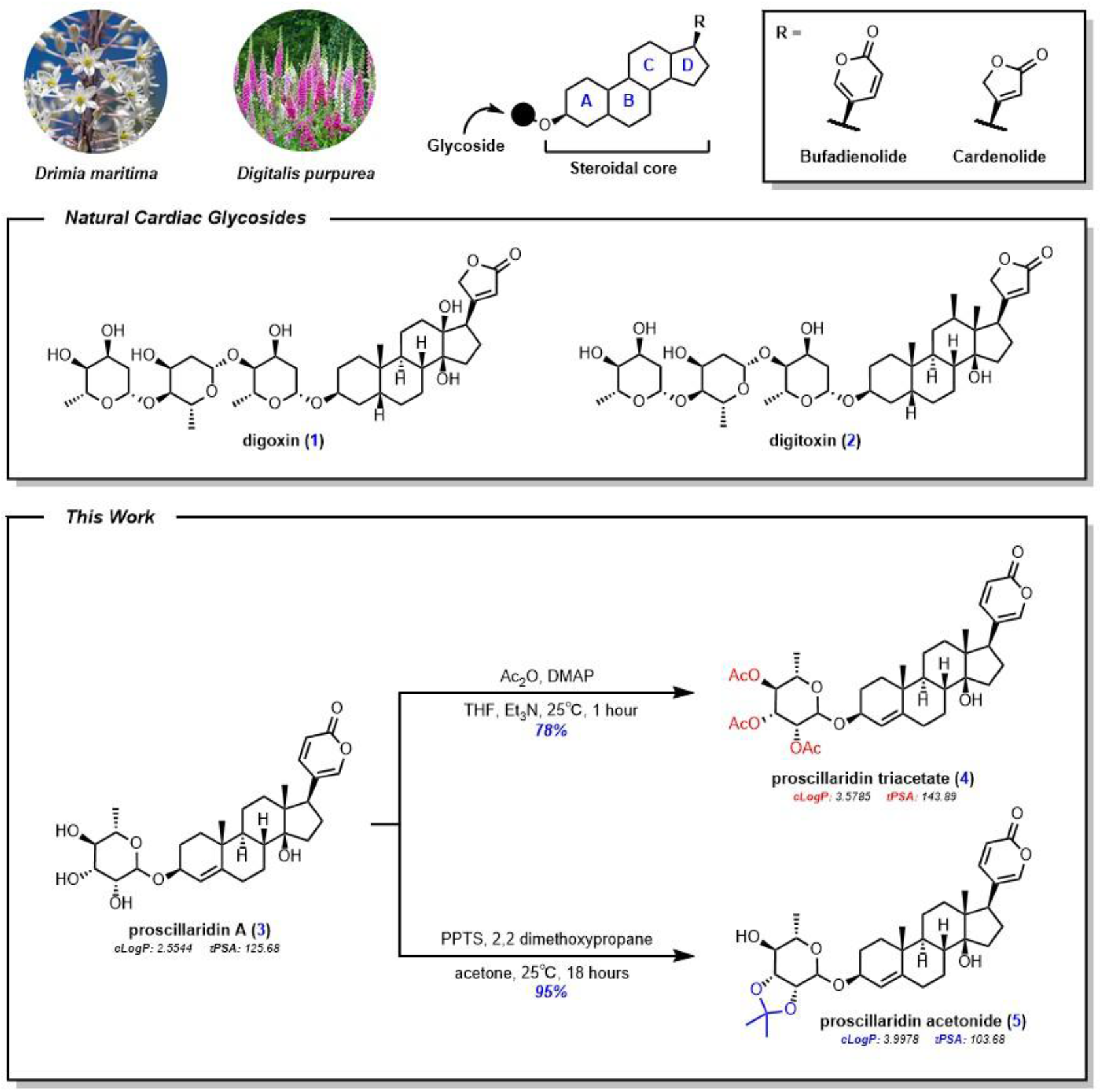
Naturally derived cardiac glycosides and their analogs have been investigated for diverse biological functions. In this work, we prepare two analogs of proscillaridin A: proscillaridin triacetate and proscillaridin acetonide. Image credits © Farmer Gracy © K. van Bourgondien

Modern clinical applications for these compounds initially focused on their potent inhibition of the Na^+^/K^+^ ATPase pump [8,9], which results in stronger heart contractions by increasing calcium levels in the sarcoplasmic reticulum. Through this mechanism, several cardiac glycosides are now being investigated as potential therapeutics for heart disease. For example, digoxin **1** has been granted full approval by the US Food and Drug Administration (FDA) for the treatment of congestive heart failure and atrial arrhythmias [10]. Additionally, digitoxin **2**, a cardenolide from the *Digitalis* genus, and proscillaridin A **3**, a bufadienolide isolated from the genus *Scilla*, have been evaluated for similar therapeutic use.

Due to their potency as plant-derived toxins, numerous cardiac glycosides have also been explored as potential anticancer agents, going beyond their primary function as cardiomyocyte modulators [1,11–14]. They have been investigated for antiproliferative activity in various cancerous cell lines, including breast cancer, osteosarcoma [15], colon cancer [16], and glioblastoma, both individually and in combination with other chemotherapeutic drugs. Several pathways have been proposed as putative targets that mediate the cytotoxic effects, including inhibition of hypoxia-inducible factor 1α (HIF-1α) [17], sensitization to TRAIL-induced cell death [18], inhibition of STAT3 activation [19], GSK-3β activation [20], targeting aberrant Myc signaling [21], and topoisomerase inhibition [22]. This suggests that this class of compounds may function through diverse intracellular mechanisms, further prompting their development as anticancer agents.

Simple modifications of other bioactive small molecules have demonstrated success in augmenting bioavailability or enhancing downstream biological activities [23–27]. In particular, ketalization [28] and esterification [29,30] have shown immense pharmacological improvements in the metabolic stability and bioavailability of compounds such as triamcinolone acetonide, a synthetic triamcinolone derivative used to treat inflammation [31], or abiraterone acetate, an FDA approved ester prodrug for the treatment of prostate cancer [32–34].

Previously, others have demonstrated that semisynthetic modification of the glycone [35], pyrone [36] or structural changes enabled by total synthesis [37] can have significant impact on anticancer potency of cardiac glycosides. Inspired by the clinical efficacy of other compounds bearing acetonides or acetate esters, we set out to investigate whether similar strategies might alter the anticancer properties of proscillaridin A **3**. Here, we synthesized two novel analogs of proscillaridin A involving acetylation **4** or ketalization **5** of the A-ring glycone. The antiproliferative activities of these compounds were evaluated alongside proscillaridin A and two additional model cardiac glycosides—digoxin **1** and digitoxin **2**—in several colorectal and liver cancer cell lines. Using a panel of *in vitro* readouts for anticancer activity in connection to pro-oncogenic and immunomodulatory pathways, including reporter cell experiments connected to TNF-α/NF-κB [38] and CD40L/NF-κB [39] pathways, IFN-γ/STAT1 [40,41], IL-10/STAT3 [42,43], IL-2/STAT5 [44,45], and Wnt1/β-catenin [46,47] pathways, we find that the ketal analog of proscillaridin A provides similar, and in some cases, enhanced *in vitro* potency.

## Results

### Chemical Synthesis

To investigate the effect of glycan peracetylation on the biological activity of proscillaridin A, we prepared the triester **4** analog in 78% yield by treatment of proscillaridin A with acetic anhydride and triethylamine. Additionally, to prepare ketal **5** from proscillaridin A, we treated the parent compound with 2,2-dimethoxypropane and catalytic pyridinium *p-*toluenesulfonate (PPTS), resulting in a 95% isolated yield of the desired glycan dimethyl ketal **5**. Compounds **4** and **5** were characterized according to conventional spectroscopic means (^1^H NMR, ^13^C NMR, FT-IR, LC-MS) and purity was established by HPLC analysis. Both the triacetate **4** and acetonide **5** exhibit much higher clogP values (3.5785 and 3.9978, respectively) compared to proscillaridin A (2.5544).

### Cell Viability Assays

In order to determine the antiproliferative activities of proscillaridin A **3** and its triacetate **4** and acetonide **5** analogs, in comparison to cardenolide cardiac glycosides digoxin **1** and digitoxin **2**, we measured cell viability at 24, 48, and 72 hours post-treatment. We observed consistent dose-dependent antiproliferative action by all three natural products and analogs **4** and **5 (Fig. 2)**. At the 24 hour timepoint, all five compounds exhibited comparable antiproliferative activity in both HCT-116 and HT-29 human colorectal cancer cell lines **(Fig. 2A, 2B)**, with IC_50_ values ranging from 0.013 μM to 1.250 μM. Notably, in CT26 murine colorectal cancer cells (which generally demonstrated lower sensitivity to all compounds compared to the three human cancer cell lines), proscillaridin A **3** and its analogs showed more potent antiproliferative activity at all three timepoints (IC_50_ values of proscillaridin A **3**: 3.887 μM, triacetate **4**: 8.088 μM, and acetonide **5**: 5.061 μM, at 72 hours) than digoxin **1** or digitoxin **2 (Fig. 2C)**. All five compounds exhibited relatively poor antiproliferative activity against HepG2 hepatocarcinoma cells at 24 hours **(Fig. 2D)**; however, all compounds showed a general increase in antiproliferative activity in HepG2 after 72 hours of treatment. In HT-29 and HCT-116 epithelial colorectal cancer cell lines, these analogs exhibited dose-dependent antiproliferative activity with significantly different IC_50_ values over all three timepoints (IC_50_ values in HT-29 and HCT-116 respectively at 72 hours for triacetate **4**: 0.132 μM and 1.230 μM, and acetonide **5**: 0.004 μM and 0.026 μM). Consistent with previous reports, proscillaridin A **3** exhibited a greater cytotoxicity than digoxin **1** and digitoxin **2**, as did its analogs in all four cell lines.

**Fig. 2.**
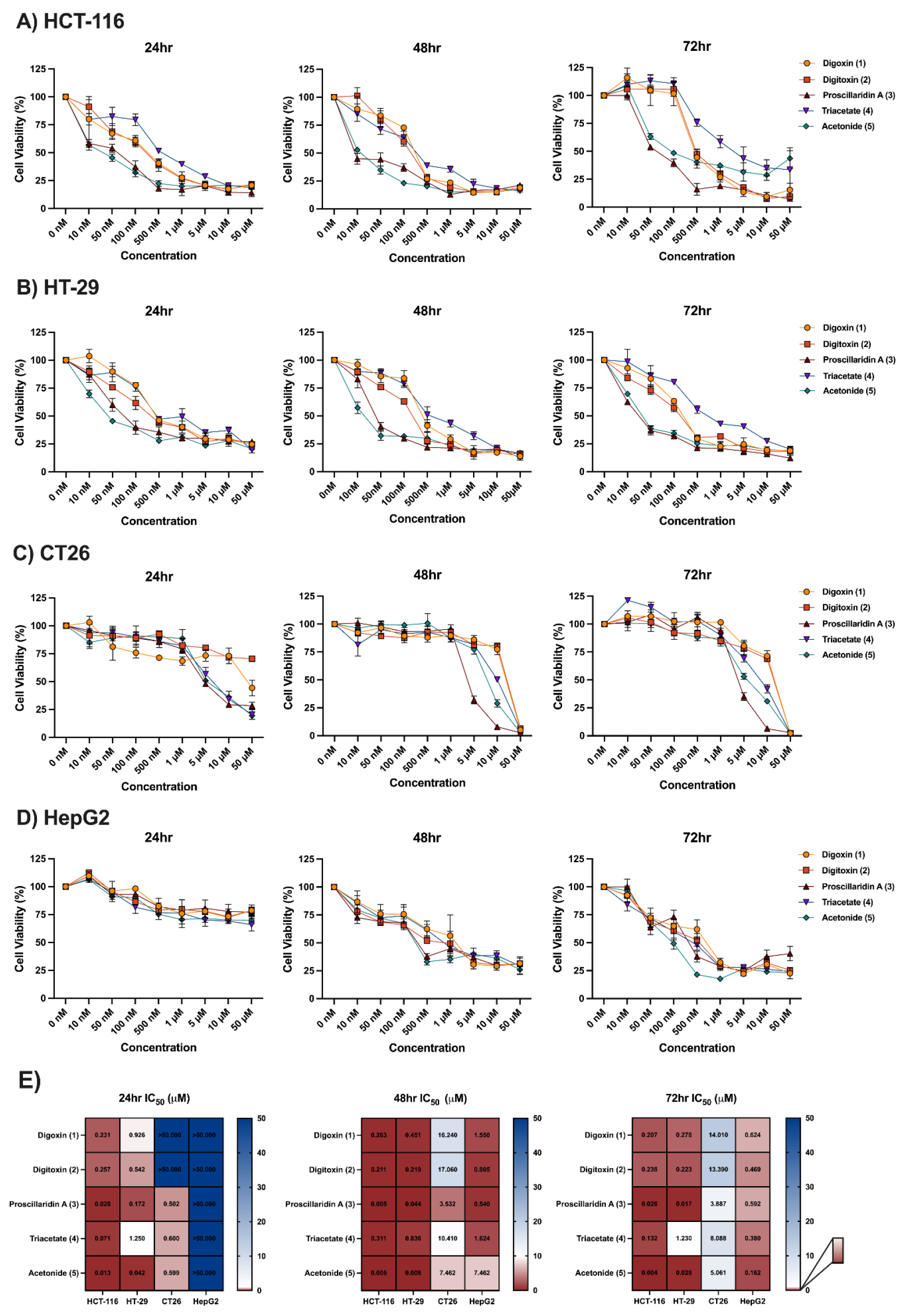
Cell viabilities of compounds evaluated in **A)** HCT-116 human colorectal cancer cells, **B)** HT-29 human colorectal cancer cells, **C)** CT26 murine colon cancer cells, and **D)** HepG2 human liver cancer cells at 24 hour, 48 hour and 72 hour timepoints. **E)** Heat map depicting cell line and time-dependent IC_50_ of cardiac glycosides and their analogs. While these compounds exhibited similar *in vitro* proliferation inhibition in HCT-116, HT-29 and CT26 cell lines, they showed strong time-dependent differences in antiproliferative action in HepG2 liver cell line. The acetonide **5** analog demonstrated an increased or similar potency to proscillaridin A **3** across all cell lines

Cell viabilities were determined by MTT (3-(4,5-dimethylthiazol-2-yl)-2,5-diphenyltetrazolium bromide) and are normalized to a control of 0.5% v/v DMSO in the absence of compound. IC_50_ values are reported as the concentration that achieves half maximal potency and were calculated on GraphPad Prism 10.4.1 **(Fig. 2E)**.

Additionally, we evaluated the cytotoxicity of compounds **1** through **5** in HCT-116 and CT26 cells by quantification of extracellular lactate dehydrogenase (LDH) activity at 72 hours according to previously reported literature protocols [48]. HCT-116 cells treated with the four highest concentrations of **1** through **5** exhibited dose-dependent increase in extracellular LDH activity, measured against a positive control of cells lysed with 9% v/v Triton X-100 and a negative control of 0.5% v/v DMSO in the absence of compound **(Fig. 3A)**. Comparatively, CT26 cells dosed at the same concentrations of **1** through **5** exhibited negligible extracellular LDH activity **(Fig. 3B)** except at the highest concentration, which is consistent with the selectivity of compounds **1** through **5** for human colon cancer cells as measured by MTT **(Fig. 2)**.

**Fig. 3.**
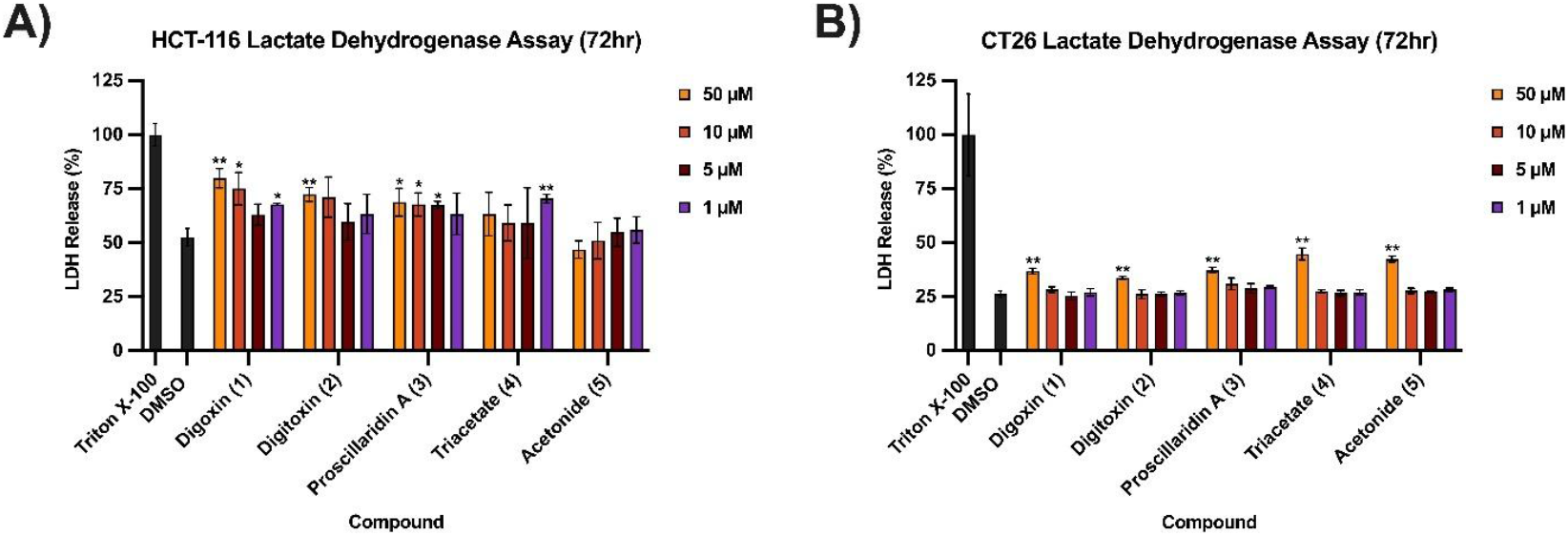
Cytotoxicity of compounds observed through lactate dehydrogenase (LDH) assays in **A)** HCT-116 human epithelial colorectal cancer cells and **B)** CT26 murine colorectal cancer cells

### Protein Marker and Cell Cycle Analysis by Flow Cytometry

Next, we measured the surface expression of several key protein markers. In order to determine if the pro-apoptotic marker caspase-3 [49,50] is implicated in the anticancer activity of our compounds, we stained HCT-116 cells treated with 100 nM or 1 μM of compounds **1** through **5** with an anti-caspase 3 primary mouse antibody labeled with a goat anti-mouse Alexa Fluor 488 conjugated secondary antibody, followed by measurement of the mean fluorescent index (MFI) via flow cytometry. We found a general increase in surface caspase-3 signal at higher doses of compounds **1** through **5**, with the most significant signal increase being in cells treated with the acetonide **5** analog **(Fig. 4A)**.

**Fig. 4.**
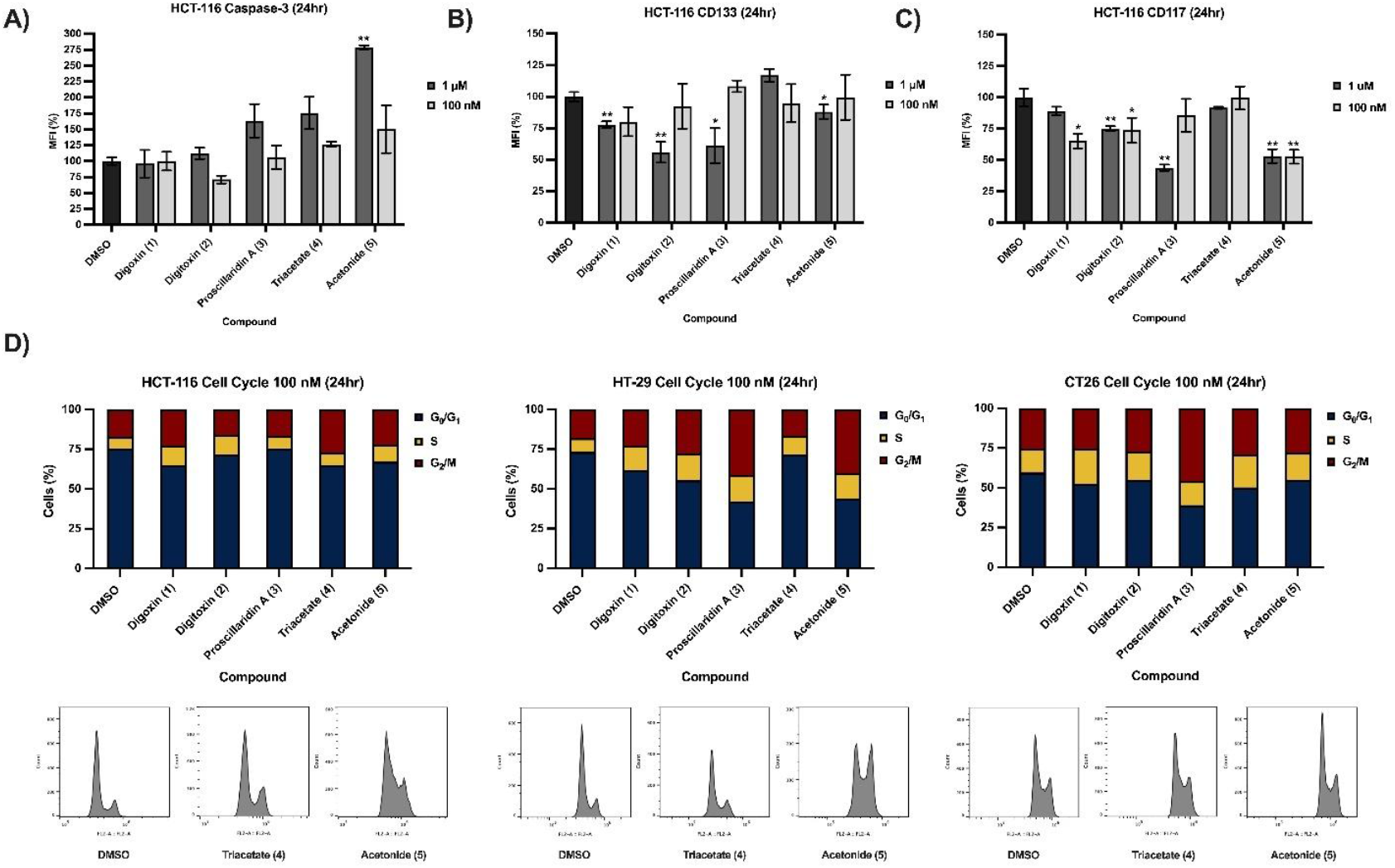
Flow cytometry profiling of key markers and cell cycle analysis in human and murine colon cancer cells. **A)** HCT-116 cells treated with 1 μM of compound **5** exhibit significantly enhanced surface expression of cleaved caspase-3, the hallmark of pre-apoptotic cell death. **B)** Effect of compounds **1** through **5** on surface CD133 (prominin 1) expression. **C)** Effect of compounds **1** through **5** on surface CD117 (c-Kit) expression. **D)** Cell cycle analysis by flow cytometry on HCT-116, HT-29, and CT26 cells treated with 100 nM of compounds **1** through **5**

Subsequently, we determined surface expression of CD133 (prominin 1), a key marker for cancer cell progression and growth in connection to epithelial mesenchymal transition (EMT) behavior in several cancers [51,52]. While minimal difference in surface CD133 expression was observed in cells treated with 100 nM of each compound, increasing the dosing of compounds **1, 2, 3** and **5** at 1 μM resulted in significant CD133 surface levels **(Fig. 4B)**.

We also sought to identify if treatment with compounds **1** through **5** affected CD117 (c-Kit) surface expression in HCT-116 cells, which is a cell growth and differentiation regulator thought to be one of the driving factors behind HCT-116 cell proliferation [53–56]. Compounds **2, 3**, and **5** resulted in significant CD117 surface level expression at 1 μM and 100 nM dosing, suggesting that CD117 might be implicated in the anticancer properties of these cardiac glycosides **(Fig. 4C)**.

Finally, to identify if cells treated with compounds **1** through **5** exhibited cell cycle arrest, we performed cell cycle analysis by flow cytometry on HCT-116 and HT-29 human colon carcinoma cells and CT26 murine colon cancer cells and observed minimal influence of compound treatment on cell cycle arrest at the 100 nM concentration **(Fig. 4D)**.

### Reporter Cell Screening

Cardiac glycosides, including proscillaridin A and its analogs, have been previously reported to putatively function through several pathways involved in cancer cell progression and immunomodulation [17– 19,21]. To evaluate the effects of treatment of cardenolides digoxin **1** and digitoxin **2** alongside proscillaridin A **3** and the triacetate **4** and ketal **5** analog, we treated several reporter cell lines with a dose titration of our compounds. In TNF-α activated NF-κB reporter cells expressing secreted extracellular alkaline phosphatase (SEAP), we observed potent dose-dependent response in cells treated with all five compounds, with the strongest inhibition of NF-κB activated SEAP activity elicited by proscillaridin A **3** and the acetonide **5** analog **(Fig. 5A)**. In a second reporter cell line with CD40L/CD40-activated NF-κB signaling for SEAP activity, we observed similar trends in potency, with the triacetate **4** analog exhibiting much weaker activity than both proscillaridin A **3** and its acetonide **5** analog **(Fig. 5B)**. Next, we subjected a STAT1/JAK reporter cell assay activated by IFN-γ to each compound and found that proscillaridin A **3** and compound **5** exhibited the most potent activity (EC_50_ = 0.033 μM and 0.067 μM respectively, **Fig. 5C**), while the triacetate **4** analog consistently returned the lowest potency in this reporter assay (EC_50_ = 2.166 μM, **Fig. 5C**). Similar trends were observed in an IL-2 activated JAK1/3-STAT5 reporter cell **(Fig. 5D)** with acetonide **5** exhibiting the most potent inhibition of STAT5-triggered SEAP activity (EC_50_ = 0.074 μM) and triacetate **4** exhibiting the lowest activity (EC_50_ = 1.933 μM). However, all compounds exhibited poor inhibition in an IL-10 activated JAK1/Tyk2-STAT3 reporter cell **(Fig. 5E)**. Remarkably, when assayed for activity in a luciferase-based Wnt1 pathway 3T3 reporter cell line with firefly luciferase (fLuc) under TCF/LEF promoter control, we observed negligible activity in cells dosed with any of the five compounds evaluated **(Fig. 5F)**. This suggests that Wnt1 signaling may not be a major contributor to the *in vitro* potency observed in such compounds. Negligible loss in cell viability was observed at the 24 hour timepoint during which endpoint analysis was performed in all reporter cell lines, strongly indicating that the dose-dependent behavior of these compounds is more closely associated with pathway interaction rather than general toxicity to the reporter cell system at hand.

**Fig. 5.**
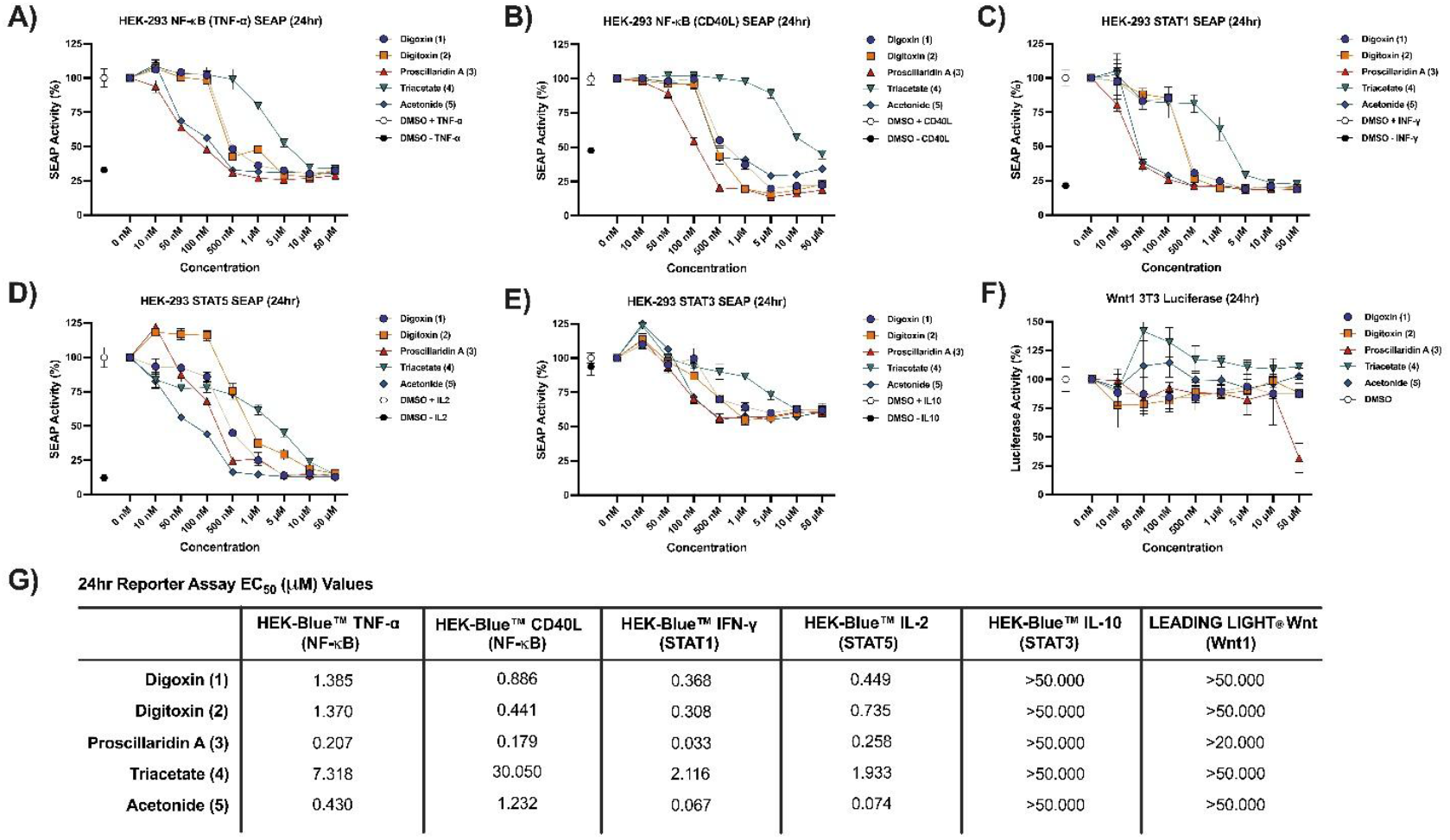
Reporter cell profiling of activity of compounds in signal pathways. **A)** SEAP activity of HEK-Blue™ TNF-α activated NF-κB reporter cells demonstrate that proscillaridin A **3** and acetonide **5** exhibit the most potent dose-dependent inhibition. **B)** SEAP activity in a CD40-activated NF-κB HEK-Blue™ reporter cell demonstrate similar profiles in NF-κB dependent SEAP activity. **C)** In HEK-Blue™ IFN-γ activated STAT1 reporter cells, proscillaridin A **3** and acetonide **5** exhibit the most potent inhibitory activity on SEAP readout. **D)** In an IL-2 activated JAK1/3-STAT5 HEK-Blue™ reporter cell, the acetonide **5** analog exhibited the most potent inhibitory activity on SEAP expression. **E)** In an IL-10 activated JAK1/Tyk2-STAT3 HEK-Blue™ reporter cell, all compounds exhibit slightly less potent inhibitory activity. **F)** In a Wnt1/TCF/LEF 3T3 luciferase-based reporter cell, all compounds exhibit minimal activity on fLuc expression. **G)** Calculated EC_50_ values of treatment of compounds **1** through **5** in a panel of six reporter cell lines

## Methods

### Synthesis Protocol

Triacetate **4** was prepared by reacting proscillaridin A **3** with acetic anhydride, triethylamine and catalytic N,N-dimethylaminopyridine (DMAP) in N,N-dimethylformamide (DMF) in 78% yield **(Fig. 1)**. Acetonide **5** was prepared by reacting proscillaridin A **3** with 2,2-dimethoxypropane and catalytic pyridinium *p-*toluenesulfonate (PPTS) in dry acetone in 95% yield **(Fig. 1)**. Spectra and detailed experimental procedures are further described in the Supporting Information.

### General Information

All reagents, solvents, and analytical standards were purchased from AK Scientific, Beantown Chemical, Acros Organics, Sigma Aldrich, HiMedia, and Cambridge Isotope Laboratories and used without further purification unless otherwise stated. McCoy’s 5A Medium, High Glucose Dulbecco’s Modified Eagle’s Medium (DMEM), RPMI-1640 Medium, 100x penicillin-streptomycin, and 0.25% trypsin EDTA were all obtained from Tribioscience (Sunnyvale, CA). Fetal bovine serum (FBS) was obtained from Gibco. Goat anti-mouse IgG(H+L)-AF488 antibody was obtained from Southern BioTech. Cleaved caspase-3 anti-mouse monoclonal antibody (Cat. #Ab156544) and CD117/c-Kit anti-mouse monoclonal antibody (Cat. #Ab096639) were obtained from Aladdin Scientific Corp. Anti-CD133 monoclonal antibody HB#7 was obtained from the DSHB and deposited there by Swaminathan, S.K./Panyam, J./Ohlfest, J.R. Ribonuclease A was obtained from Research Products International (97% purity).

### Cell Culture

HCT-116 and HT-29 human colorectal cancer cell lines were obtained from the European Collection of Authenticated Cell Cultures (ECACC) and cultured in McCoy’s 5A Medium (Tribioscience) supplemented with 10% v/v fetal bovine serum (FBS, Gibco) and 1% v/v 100x penicillin-streptomycin (Tribioscience). CT26, a murine colorectal carcinoma cell line, was obtained from American Type Culture Collection (ATCC), and cultured in RPMI-1640 Medium (Tribioscience) supplemented with 10% v/v fetal bovine serum (FBS, Gibco) and 1% v/v 100x penicillin-streptomycin (Tribioscience). HepG2, a human liver cancer cell line, was obtained from the European Collection of Authenticated Cell Cultures (ECACC) and cultured in High Glucose Dulbecco’s Modified Eagle Medium (DMEM) (Hygia Reagents) supplemented with 10% v/v fetal bovine serum (FBS, Gibco) and 1% v/v 100x penicillin-streptomycin (Tribioscience). A human embryonic kidney cell line, HEK-Blue™ TNF-α, stably transfected with a TNF-α inducible secreted alkaline phosphatase (SEAP) reporter gene, was obtained from Invivogen (Cat. #hkb-tnfdmyd) and was cultured in High Glucose Dulbecco’s Modified Eagle Medium (DMEM) (Hygia Reagents) supplemented with 10% v/v fetal bovine serum (FBS, Gibco) and 1% v/v 100x penicillin-streptomycin (Tribioscience). HEK-Blue™ CD40L cells engineered to express the CD40 receptor were obtained from Invivogen (Cat. #hkb-cd40) and cultured in High Glucose Dulbecco’s Modified Eagle Medium (DMEM) (Hygia Reagents), supplemented with 10% v/v fetal bovine serum (FBS, Gibco) and 1% v/v 100x penicillin-streptomycin (Tribioscience). HEK-Blue™ IFN-γ cells engineered to monitor human type II interferon IFN-γ-induced STAT1 stimulation were obtained from Invivogen (Cat. #hkb-ifng) and cultured in High Glucose Dulbecco’s Modified Eagle Medium (DMEM) (Hygia Reagents), supplemented with 10% v/v fetal bovine serum (FBS, Gibco) and 1% v/v 100x penicillin-streptomycin (Tribioscience). HEK-Blue™ IL-10 cells engineered for the detection of human IL-10 to monitor JAK1/STAT3 signaling were obtained from Invivogen (Cat. #hkb-il10) and cultured in High Glucose Dulbecco’s Modified Eagle Medium (DMEM) (Hygia Reagents), supplemented with 10% v/v fetal bovine serum (FBS, Gibco) and 1% v/v 100x penicillin-streptomycin (Tribioscience). HEK-Blue™ IL-2 cells engineered for the detection of human IL-2 to monitor JAK1/3-STAT5 signaling were obtained from Invivogen (Cat. #hkb-il2) and cultured in High Glucose Dulbecco’s Modified Eagle Medium (DMEM) (Hygia Reagents), supplemented with 10% v/v fetal bovine serum (FBS, Gibco) and 1% v/v 100x penicillin-streptomycin (Tribioscience). LEADING LIGHT® Wnt Reporter 3T3 mouse embryonic fibroblast cells were obtained from Enzo Life Sciences (Cat. #ENZ-61002-0001) and cultured in High Glucose Dulbecco’s Modified Eagle Medium (DMEM) (Hygia Reagents), supplemented with 10% v/v fetal bovine serum (FBS, Gibco), and 1% v/v 100x penicillin-streptomycin (Tribioscience).

HCT-116, HT-29, CT26, HepG2, HEK-293, and 3T3 cells were cultured in T25 and T75 flasks (Corning), in a 37°C incubator (5.0% CO_2_). Cultures were maintained by splitting cells 1:4 at 70% confluency approximately every 3 days.

### Cell Viability Assays

#### MTT Assay

Cell cultures were grown according to the procedure above. To plate cells, a confluent culture was detached from a T75 flask (Corning) using 0.25% trypsin-EDTA, resuspended in media and plated in a 96-well flat bottom tissue culture treated plate (Corning) with 100 µL per well. Following a 24 hour incubation period at 37°C (5.0% CO_2_), a drug medium solution was prepared by adding 200-fold compound solutions dissolved in dimethyl sulfoxide (DMSO) to cell culture media. Old media was aspirated and replaced with 110 µL, 140 µL and 170 µL of fresh media–for 24-, 48- and 72-hour time intervals, respectively–of drug medium solution in quadruplicate. A negative control of 0.5% v/v DMSO was also included. The plates were then incubated at 37°C (5.0% CO_2_) for 24, 48, and 72 hours, after which 10 µL of a fresh solution of 3-(4,5-dimethylthiazol-2-yl)-2,5-diphenyltetrazolium bromide (MTT) (AK Scientific) in 1x phosphate-buffered saline (PBS) (5 mg/mL) was added to all wells and mixed. Plates were incubated at 37°C (5.0% CO_2_) for an additional 2-4 hours before aspiration and the addition of 100 µL of DMSO, which was mixed until all formazan crystals were fully dissolved. Absorbance was measured with a Molecular Devices SPECTRAmax 250 Microplate Spectrophotometer at 570 nm. Cell viability was calculated from the average normalized absorbance of each treatment group and plotted against drug concentration. A logarithmic trend line was used to determine IC_50_ values by solving for the x-coordinate (drug concentration) when y (percent cell viability) reached exactly 50%. IC_50_ values were determined using GraphPad Prism 10.4.1.

#### Lactate Dehydrogenase (LDH) Assay

Extracellular LDH activity, released due to a loss of cell membrane integrity, was measured using a previously reported Cold Spring Harbor Protocol. HCT-116 and CT26 cells were seeded at 70% confluency in McCoy’s 5A Medium and RPMI-1640 respectively, supplemented with 10% v/v fetal bovine serum (FBS, Gibco) and 1% v/v 100x penicillin-streptomycin (Tribioscience), and treated with 200-fold dilutions of compounds **1** through **5** to reach final concentrations of 50 µM, 10 µM, 5 µM, 1 µM, 500 nM, 100 nM, 50 nM and 10 nM. A negative control of 0.5% v/v DMSO was also included. At 72 hours, to a negative control of cells drugged with DMSO was added 15 μL of lysis solution (9% v/v Triton X-100) for 5 minutes to act as a positive control for maximum LDH release. Then, 50 µL of cell media from each treatment was moved to a non-tissue culture treated 96-well plate and allowed to react with 50 µL of the LDH substrate solution (L-(+)-lactic acid (0.054 M), β-NAD^+^ (1.30 mM), 1-Methoxy-5-methylphenazinium methyl sulfate (0.28 mM), and 2-*p*-iodophenyl-3-*p*-nitrophenyl tetrazolium chloride (INT) solution (0.66 mM) dissolved in 0.2 M Tris-HCl buffer (pH 8.2)), for 45 min at 37°C (5.0% CO_2_) protected from light before the addition of 100 µL of stop solution solution (50% dimethylformamide and 20% SDS at pH 4.7) to each sample well. The final product was measured spectrophotometrically at 490 nm and 750 nm using a Molecular Devices SPECTRAmax 250 Microplate Spectrophotometer. The absorbance taken at 750 nm was subtracted from the 490 nm absorbance value to calculate LDH activity.

### Flow Cytometry

#### Cell Cycle Analysis

HCT-116, HT-29, and CT26 cells were seeded at 70% confluency in 6-well tissue-culture treated plates (Corning) in 1.50 mL of cell-line respective media (McCoy’s 5A Medium for HCT-116 and HT-29, RPMI-1640 for CT26) supplemented with 10% v/v fetal bovine serum (FBS, Gibco) and 1% v/v 100x penicillin-streptomycin (Tribioscience). Cells were incubated for 24 hours at 37°C (5.0% CO_2_) to allow for attachment, after which they were treated with 100 nM of compounds **1** through **5**. A negative control of 0.5% v/v DMSO was also included. After 24 hours of incubation with the drug, the supernatant media was collected in 2 mL Eppendorf tubes and the cells were trypsinized with 400 µL of 0.25% trypsin-EDTA (Tribioscience) per well. The cells were incubated for 10 minutes, and the trypsin was deactivated with 300 µL of cell-line respective media. The detached cells were moved into the same Eppendorf tubes as those containing their drugged media and centrifuged for 9 minutes at 2500 rpm. The supernatant was removed, the cell pellet was washed in 500 µL of 1x PBS and re-centrifuged for 10 minutes at 3500 rpm. The PBS was aspirated off and the cells were resuspended in 500 µL of 70% ethanol at 4°C for 30 minutes, then re-pelleted by centrifugation. The ethanol was aspirated off, the cells were resuspended in 25 µL of ribonuclease A (1 mg/mL in DI water) and 100 µL of PBS and were then incubated for 5 minutes at 37°C (5.0% CO_2_). The cells were then stained in the dark with 1 µL propidium iodide (1 mg/mL in DI water, AK Scientific, 98%) for 30 minutes. DNA content was then analyzed to determine cell cycle stage with a BD Accuri C6 Flow Cytometer. Each sample was run to 50,000 events and single cells were gated to exclude debris, based on forward scatter and side scatter, followed by FL2-A channel gating. Results were analyzed using FlowJo Software 10 to determine the percentage of cells in each stage of the cell cycle (G_0_/G_1_, S, G_2_/M).

#### Caspase-3 Flow Cytometry

HCT-116 cells were seeded at 80% confluency in 12-well tissue-culture treated plates (Corning) with McCoy’s 5A Medium supplemented with 10% v/v fetal bovine serum (FBS, Gibco) and 1% v/v 100x penicillin-streptomycin (Tribioscience). Cells were incubated for 24 hours at 37°C (5.0% CO_2_) to allow for attachment, after which they were treated with 100 nM and 1 µM of compounds **1** through **5**. A negative control of 0.5% v/v DMSO was also included. After 24 hours of incubation with the drug, the supernatant media was collected in 2 mL Eppendorf tubes and the cells were trypsinized with 300 µL of 0.25% trypsin-EDTA (Tribioscience) per well. The cells were incubated for 10 minutes, and the trypsin was deactivated with 200 µL of McCoy’s 5A Medium. The detached cells were transferred into the same Eppendorf tubes as their drugged media and centrifuged for 9 minutes at 2500 rpm. The supernatant was removed, the cell pellet was washed in 300 µL of 1x phosphate-buffered saline (PBS) and re-centrifuged for 10 minutes at 3500 rpm. The PBS was then aspirated off and the cells were resuspended in 100 μL of 1.0% bovine serum albumin (BSA) and 1 μL of cleaved caspase-3 anti-mouse mAb (300 μg/ml in 1.0% BSA). The cells were incubated in the dark at 4°C for 1 hour, re-pelleted by centrifugation and washed with 500 µL of 1x PBS. The PBS was removed, and the cells were resuspended in 100 μL of 1.0% BSA and 0.40 μL of goat anti-mouse IgG(H+L)-AF488 antibody (1 mg/mL). The samples were incubated once more in the dark at 4°C for 1 hour before mean fluorescent index (MFI) was measured with the BD Accuri C6 Flow Cytometer. Each sample was run to 50,000 events and single cells were gated to exclude debris, based on forward scatter and side scatter. MFI was calculated against FL1-A and expressed as percentages in relation to the control. Results were analyzed using FlowJo Software 10.

#### CD117 Flow Cytometry

HCT-116 cells were seeded at 80% confluency in 6-well tissue-culture treated plates (Corning) with McCoy’s 5A Medium supplemented with 10% v/v fetal bovine serum (FBS, Gibco) and 1% v/v 100x penicillin-streptomycin (Tribioscience). Cells were incubated for 24 hours at 37°C (5.0% CO_2_) to allow for attachment, after which they were treated with 100 nM and 1 µM of compounds **1** through **5**. A negative control of 0.5% v/v DMSO was also included. After 24 hours of incubation with the drug, the supernatant media was collected in 2 mL Eppendorf tubes and the cells were trypsinized with 300 µL of 0.25% trypsin-EDTA (Tribioscience) per well. The cells were incubated for 10 minutes, and the trypsin was deactivated with 200 µL of McCoy’s 5A Medium. The detached cells were transferred into the same Eppendorf tubes as their drugged media and centrifuged for 9 minutes at 2500 rpm. The supernatant was removed, the cell pellet was washed in 300 µL of 1x phosphate-buffered saline (PBS) and re-centrifuged for 10 minutes at 3500 rpm. The PBS was then aspirated off and the cells were resuspended in 100 μL of 1.0% bovine serum albumin (BSA) and 1 μL of CD117 anti-mouse mAb (300 μg/ml in 1.0% BSA). The cells were incubated in the dark at 4°C for 1 hour, re-pelleted by centrifugation and washed with 500 µL of 1x PBS. The PBS was removed, and the cells were resuspended in 100 μL of 1.0% BSA and 0.40 μL of goat anti-mouse IgG(H+L)-AF488 antibody (1 mg/mL). The samples were incubated once more in the dark at 4°C for 1 hour before mean fluorescent index (MFI) was measured with the BD Accuri C6 Flow Cytometer. Each sample was run to 50,000 events and single cells were gated to exclude debris, based on forward scatter and side scatter. MFI was calculated against FL1-A and expressed as percentages in relation to the control. Results were analyzed using FlowJo Software 10.

#### CD133 Flow Cytometry

HCT-116 cells were seeded at 80% confluency in 12-well tissue-culture treated plates (Corning) with 800 µL of McCoy’s 5A Medium supplemented with 10% v/v fetal bovine serum (FBS, Gibco) and 1% v/v 100x penicillin-streptomycin (Tribioscience). Cells were incubated for 24 hours at 37°C (5.0% CO_2_) to allow for attachment, after which they were treated with 100 nM and 1 µM of compounds **1** through **5**. A negative control of 0.5% v/v DMSO was also included. After 24 hours of incubation with the drug, the supernatant media was collected in 2 mL Eppendorf tubes and the cells were trypsinized with 300 µL of 0.25% trypsin-EDTA (Tribioscience) per well. The cells were incubated for 10 minutes, and the trypsin was deactivated with 200 µL of McCoy’s 5A Medium. The detached cells were moved into the same Eppendorf tubes as their drugged media and centrifuged for 9 minutes at 2500 rpm. The supernatant was removed, the cell pellet was washed in 300 µL of 1x phosphate-buffered saline (PBS) and re-centrifuged for 10 minutes at 3500 rpm. The PBS was then aspirated off and the cells were resuspended in 100 μL of 1.0% bovine serum albumin (BSA) and 9.09 μL of CD133 anti-mouse mAb (36 μg/ml in 1.0% BSA). The cells were then incubated in the dark at 4°C for 1 hour, re-pelleted by centrifugation and washed with 500 µL of 1x PBS. The PBS was removed, and the cells were resuspended in 100 μL of 1.0% BSA and 0.40 μL of goat anti-mouse IgG(H+L)-AF488 antibody (1 mg/mL). The samples were incubated once more in the dark at 4°C for 1 hour before mean fluorescent index (MFI) was measured with the BD Accuri C6 Flow Cytometer. Each sample was run to 50,000 events and single cells were gated to exclude debris based on forward scatter and side scatter. MFI was calculated against FL1-A and expressed as percentages in relation to the control. Results were analyzed using FlowJo Software 10.

### Reporter Assays

#### NF-κB (TNF-α) Reporter Secreted Alkaline Phosphatase Cell Assay

HEK-Blue™ TNF-α cells, obtained from Invivogen (Cat. #hkd-tnfa), were seeded at 80% confluency in a 96-well tissue culture treated flat bottom plate (Corning) with 100 µL per well and incubated at 37°C (5.0% CO_2_) for 24 hours. Seeding media was then aspirated off and a drug solution prepared using TNF-α laced (2 ng/mL, Tribioscience) High Glucose Dulbecco’s Modified Eagle’s Medium was added reaching final concentrations of compounds **1** through **5** ranging from 50 µM to 10 nM. A negative control of 0.5% v/v DMSO with TNF-α laced media and with regular media was also included. The general protocol for preparing the para-Nitrophenylphosphate solution is as follows: the para-Nitrophenylphosphate solution was prepared to final concentrations of 10% v/v diethanolamine, 0.10 mM magnesium chloride, 5 mg/mL of para-Nitrophenylphosphate and adjusted to a pH of 9.8 in DI water. Following 24 hours incubation at 37°C (5.0% CO_2_), 10 µL of culture supernatant from SEAP-expressing cells was added to 90 µL of a para-Nitrophenylphosphate solution (prepared using the general protocol) in a secondary 96-well plate. After incubation for 45 minutes at 37°C (5.0% CO_2_), optical density was measured at 405 nm using a Molecular Devices SPECTRAmax 250 Microplate Spectrophotometer. EC_50_, or the half-maximal effective concentration value, is reported as the concentration of the drug required to produce a 50% response and was calculated on GraphPad Prism 10.4.1.

#### NF-κB (CD40L) Reporter Secreted Alkaline Phosphatase Cell Assay

HEK-Blue™ CD40L cells, obtained from Invivogen (Cat. #hkb-cd40), were seeded at 70% confluency in a 96-well tissue culture treated flat bottom plate (Corning) with 100 µL per well and incubated at 37°C (5.0% CO_2_) for 24 hours. Seeding media was then aspirated off and a drug solution prepared with CD40L laced (2 ng/mL, Tribioscience) High Glucose Dulbecco’s Modified Eagle’s Medium was added reaching final concentrations of compounds **1** through **5** ranging from 50 µM to 10 nM. A negative control of 0.5% v/v DMSO with CD40L laced media and with regular media was also included. Following 24 hours incubation at 37°C (5.0% CO_2_), 10 µL of culture supernatant from SEAP-expressing cells was added to 90 µL of a para-Nitrophenylphosphate solution (prepared using the general protocol) in a secondary 96-well plate. The optical density was measured immediately at 405 nm using a Molecular Devices SPECTRAmax 250 Microplate Spectrophotometer. EC_50_ values were calculated on GraphPad Prism 10.4.1.

#### STAT1 Reporter Secreted Alkaline Phosphatase Cell Assay

HEK-Blue™ IFN-γ cells, obtained from Invivogen (Cat. #hkb-ifng) were seeded at 70% confluency in a 96-well tissue culture treated flat bottom plate (Corning) with 100 µL per well and incubated at 37°C (5.0% CO_2_) for 24 hours. Seeding media was then aspirated off and a drug medium solution prepared with IFN-γ laced (2 ng/mL, Tribioscience) High Glucose Dulbecco’s Modified Eagle’s Medium was added reaching final concentrations of compounds **1** through **5** ranging from 50 µM to 10 nM. A negative control of 0.5% v/v DMSO with IFN-γ laced media and with regular media was also included. Following 24 hours incubation at 37°C (5.0% CO_2_), 10 µL of culture supernatant from SEAP-expressing cells was added to 90 µL of a para-Nitrophenylphosphate solution (prepared using the general protocol) in a secondary 96-well plate. After incubation for 15 minutes at 37°C (5.0% CO_2_), optical density was measured at 405 nm using a Molecular Devices SPECTRAmax 250 Microplate Spectrophotometer. EC_50_ values were calculated on GraphPad Prism 10.4.1.

#### STAT5 Reporter Secreted Alkaline Phosphatase Cell Assay

HEK-Blue™ IL-2 cells, obtained from Invivogen (Cat. #hkb-il2) were seeded at 70% confluency in a 96-well tissue culture treated flat bottom plate (Corning) with 100 µL per well and incubated at 37°C (5.0% CO_2_) for 24 hours. Seeding media was then aspirated off and a drug solution prepared with IL-2 laced (1 ng/mL, Tribioscience) High Glucose Dulbecco’s Modified Eagle’s Medium was added reaching final concentrations of compounds **1** through **5** ranging from 50 µM to 10 nM. A negative control of 0.5% v/v DMSO with IL-2 laced media and with regular media was also included. Following 24 hours incubation at 37°C (5.0% CO_2_), 10 µL of culture supernatant from SEAP-expressing cells was added to 90 µL of a para-Nitrophenylphosphate solution (prepared using the general protocol) in a secondary 96-well plate. The optical density was measured immediately at 405 nm using a Molecular Devices SPECTRAmax 250 Microplate Spectrophotometer. EC_50_ values were calculated on GraphPad Prism 10.4.1.

#### STAT3 Reporter Secreted Alkaline Phosphatase Cell Assay

HEK-Blue™ IL-10 cells, obtained from Invivogen (Cat. #hkb-il10) were seeded at 70% confluency in a 96-well tissue culture treated flat bottom plate (Corning) with 100 µL per well and incubated at 37°C (5.0% CO_2_) for 24 hours. Seeding media was then aspirated off and a drug solution prepared with IL-10 laced (1 ng/mL, Tribioscience) High Glucose Dulbecco’s Modified Eagle’s Medium was added reaching final concentrations of compounds **1** through **5** ranging from 50 µM to 10 nM. A negative control of 0.5% v/v DMSO with IL-10 laced media and with regular media was also included. Following 24 hours incubation at 37°C (5.0% CO_2_), 10 µL of culture supernatant from SEAP-expressing cells was added to 90 µL of a para-Nitrophenylphosphate solution (prepared using the general protocol) in a secondary 96-well plate. After incubation for 20 minutes at 37°C (5.0% CO_2_), optical density was measured at 405 nm using a Molecular Devices SPECTRAmax 250 Microplate Spectrophotometer. EC_50_ values were calculated on GraphPad Prism 10.4.1.

#### Wnt1 Luciferase Reporter Cell Assay

LEADING LIGHT® Wnt Reporter 3T3 mouse embryonic fibroblast cells were obtained from Enzo LifeSciences (Cat. #ENZ-61002-0001) and seeded at 80% confluency in 96-well flat bottom plates (Corning). After 24 hours of incubation at 37°C (5.0% CO_2_), the plates were treated for 24 hours with compounds **1** through **5** at final concentrations ranging from 50 µM to 10 nM. The overexpression of β-catenin/TCF/LEF signaling factors was induced by the coadministration of 10 µM CHIR-99021 (AK Scientific). A negative control of 0.5% v/v DMSO and a background control were used to establish background signal intensity in the absence of an inhibitor, and background signal in the absence of luciferin chemiluminescence. After 24 hours of incubation at 37°C (5.0% CO_2_), 70 μL of the culture supernatant was transferred to a black wall, flat bottom opaque 96-well plate (Corning) and 70 μL of the 3x Firefly Assay Buffer (15 mM dithiothreitol, 0.60 mM coenzyme A, 0.45 mM ATP, 4.2 mg/mL D-luciferin, Triton X-100 lysis buffer) was added on top. Luminescence was quantified using a Tecan FarCyte Ultra Plate Reader. EC_50_ values were calculated on GraphPad Prism 10.4.1.

#### Calculation of tPSA and cLogP

For our analogs, the tPSA values, which measure the total polar surface area for a molecule, and cLogP values, which represent the calculated lipophilicity for a compound, were calculated using ChemDraw Professional 22.2.0.

### Statistical Analysis

All statistical analyses were performed using GraphPad Prism 10.4.1. Data in each group were compared using a Welch’s t-test. P-values 0.05 were considered statistically significant.

## Conclusions

Inspired by two common strategies to enhance the biological activity of bioactive small molecules through chemical modification, we prepared two novel analogs of the anticancer cardiac glycoside proscillaridin A, including peracetylation of the A-ring glycan to afford triacetate **4** and installation of a dimethyl ketal on the A-ring glycan to afford acetonide **5**. We evaluated the antiproliferative activity of these two synthetic proscillaridin A analogs in comparison to the natural product and digoxin **1** and digitoxin **2**, two other cardiac glycosides bearing a cardenolide lactone ring, and found that acetonide **5** exhibited potent action as an antiproliferative agent in epithelial and murine colorectal cancer cells. All three cardiac glycoside natural products and their analogs exhibited cytotoxic behavior in human colorectal cancer cells, while potency was generally attenuated in CT26 murine colon cancer cells and HepG2 human hepatocarcinoma cells. In HepG2 cells, we observed a clear increase in antiproliferative activity with increasing timepoints, while potent inhibition of colorectal cancer cells was observed even after only 24 hours of treatment with compounds. Additionally, we observed minimal influence of compound treatment on cell cycle arrest through flow cytometry. Further, we profiled their activity in altering surface expression of key markers, including CD117 (c-Kit), CD133 (prominin 1), and cleaved caspase-3. We found that all three cardiac glycoside natural products and acetonide **5** analog inhibit surface expression of the pro-cancer markers CD117 and CD133 while acetonide **5** significantly increases the presence of surface-marked caspase-3, shedding light into potential signaling pathways through which these compounds exert their anticancer activity.

Finally, we evaluated the activity of the three cardiac glycosides alongside our two novel analogs in a panel of six reporter cell assays, illuminating several key cellular pathways involved in cell survival and proliferation, tumorigenesis, and immunomodulation, including those connected to TNF-α/NF-κB, CD40/NF-Kb. Wnt1/β-catenin, IFN-γ/STAT1, IL-2/STAT5, and IL-10/STAT3 signaling. We observed potent inhibition of NF-κB activated expression of secreted alkaline phosphatase (SEAP) activity in both TNF-α and CD40-triggered HEK-Blue™ reporter cells, with the most potent activity observed in cells treated with proscillaridin A **3** and its acetonide **5** analog. Moreover, in all three STAT-dependent HEK-Blue™ reporter cells, including an INF-γ activated STAT1 reporter, IL-2 activated STAT5 reporter, and an IL-10 activated STAT3 reporter, we observed similar dose-dependent trends with decreasing SEAP activity, albeit with attenuated potency in the IL-10 activated STAT3 reporter cells. Remarkably, in a Wnt1 3T3 reporter cell expressing firefly luciferase (fLuc) under TCF/LEF control, we observed minimal activity of all compounds, even at the highest doses.

Collectively, these results suggest that certain synthetic glycan modifications are tolerated in retaining the biological potency of proscillaridin A **3** as an anticancer agent. We demonstrate that this is broadly applicable in a panel of colorectal and liver cancer cell lines. These initial findings prompt further development of glycan modified analogs of proscillaridin A, as well as *in vivo* studies to assess the bioavailability and metabolic stability of these and other modified cardiac glycosides. Such studies are now underway in our laboratories.

## Author Contributions

S.S., L.M., and E.N. performed the biological assays, including MTT assays, reporter cell assays, and flow cytometry. L.M. and S.S. performed data analysis and prepared figures. S.H. and E.N. synthesized the compounds used in this study. F.W.J., G.J., and E.N. conceived the project and supervised the study. S.S., L.M., F.W.J., G.J., and E.N. wrote the manuscript.

## Acknowledgements

The authors gratefully acknowledge Dr. Stephen Lynch at the Stanford University Nuclear Magnetic Resonance (NMR) facility for access to high field NMR spectra for the novel analogs reported in this paper. Additionally, the authors gratefully acknowledge Dr. Zhong Wang for generously providing HEK-Blue™ reporter cell lines and Dr. Darrin Liang (Tribioscience, Sunnyvale, CA) for generously donating IL-10 and other bioreagents employed in this study. Data analysis was performed on Prism GraphPad Prism 10.4.1, spectroscopic (NMR) data was analyzed on MestreNova™, flow cytometry data was processed on FlowJo Software 10, and figures were generated on BioRender™ or Revvity Signals ChemDraw™.

