## Supporting Information Document for "Synthesis, anticancer properties, and biological profiling of synthetic glycan analogs of proscillaridin A"

### CONTENTS


1. General Information
    a. Materials SI-3
    b. Equipment SI-3

2. Experimental Procedures SI-4
    a. Chemical Synthesis & Characterization SI-4
        i. proscillaridin triacetate SI-4
        ii. proscillaridin acetonide SI-8
    b. Cell Culture SI-12
    c. Cell Viability Assays SI-13
        i. MTT Assay SI-13
        ii. Lactate Dehydrogenase Assay SI-13
    d. Flow Cytometry SI-14
        i. Cell Cycle Analysis SI-14
        ii. Caspase-3 SI-14
        iii. CD117 SI-15
        iv. CD133 SI-15
    e. Reporter Cell Assays SI-16
        i. NF-κB (TNF-α) Reporter Secreted Alkaline Phosphatase Cell Assay SI-16
        ii. NF-κB (CD40L) Reporter Secreted Alkaline Phosphatase Cell Assay SI-16
        iii. STAT1 Reporter Secreted Alkaline Phosphatase Cell Assay SI-16
        iv. STAT5 Reporter Secreted Alkaline Phosphatase Cell Assay SI-17
        v. STAT3 Reporter Secreted Alkaline Phosphatase Cell Assay SI-17
        vi. Wnt1 Luciferase Reporter Cell Assay SI-17

3. Supplementary Figures SI-19
    A. NF-κB (TNF-α) Reporter Secreted Alkaline Phosphatase Cell Assay SI-19
    B. NF-κB (CD40L) Reporter Secreted Alkaline Phosphatase Cell Assay SI-19
    C. STAT1 Reporter Secreted Alkaline Phosphatase Cell Assay SI-19
    D. STAT5 Reporter Secreted Alkaline Phosphatase Cell Assay SI-19
    E. STAT3 Reporter Secreted Alkaline Phosphatase Cell Assay SI-19
    F. Wnt1 Luciferase Reporter Cell Assay SI-19

### 1. General Information

#### a. Materials:

Solvents used in all reactions and purification processes were ACS grade or higher and were used without additional purification. They were purchased from Fisher Chemical, Sigma Aldrich, Sierra Chemical Corp, Beantown Chemical, Stellar Chemical, JT Baker, or Acros Organics. Deuterated solvents were purchased from Cambridge Isotope Laboratories or Acros Organics and were used without further purification. Solvents used in analytical methods (HPLC, LCMS) were HPLC grade (22 micron filtered). All reagents, catalysts, and chemicals were purchased from commercial sources and used without further purification unless otherwise stated.

#### b. Equipment:

$^{1}$H and $^{13}$C{$^{1}$H} NMR spectra were acquired on a Varian INOVA 400 MHz nuclear magnetic resonance spectrometer, a Bruker Avance Neo 400 MHz nuclear magnetic resonance spectrometer, or Nanalysis NMReady 60Pro multinuclear benchtop nuclear magnetic resonance spectrometer and were processed on the MestreNova software package. $^{1}$H and $^{13}$C chemical shifts are reported in parts per million (ppm). $^{1}$H chemical shifts are reported relative to the residual solvent peak ($CDCl_3$ = 7.26 ppm) as follows: chemical shift (δ), multiplicity (app = apparent, b = broad, s = singlet, d = doublet, t = triplet, q = quartet, m = multiplet, or combinations thereof), coupling constant(s) in Hz, integration. $^{13}$C chemical shifts are reported relative to the residual solvent peak ($CDCl_3$ = 77.00 ppm). Mass spectra were obtained using a Thermo Electron LTQ-XL linear ion trap mass spectrometer equipped with a Thermo Finnigan Surveyor reverse phase high performance liquid chromatography (LC-MS) or a Waters Micromass Quattro triple quadrupole mass spectrometer. Infrared spectra were collected on a Thermo Scientific Nicolet iS5 Fourier transform infrared (FT-IR) spectrometer equipped with a Thermo iD5 attenuated total reflectance (ATR) assembly. MTT, LDH, and SEAP data were acquired on a Molecular Devices SPECTRAmax 250 Microplate Spectrophotometer. Luminescence data was acquired on a Molecular Devices SPECTRAmax Paradigm. Flow cytometry data was acquired on a BD Accuri C6 flow cytometer.

#### 2. Experimental Procedures

##### a. Chemical Synthesis & Characterization

**Preparation of proscillaridin triacetate**

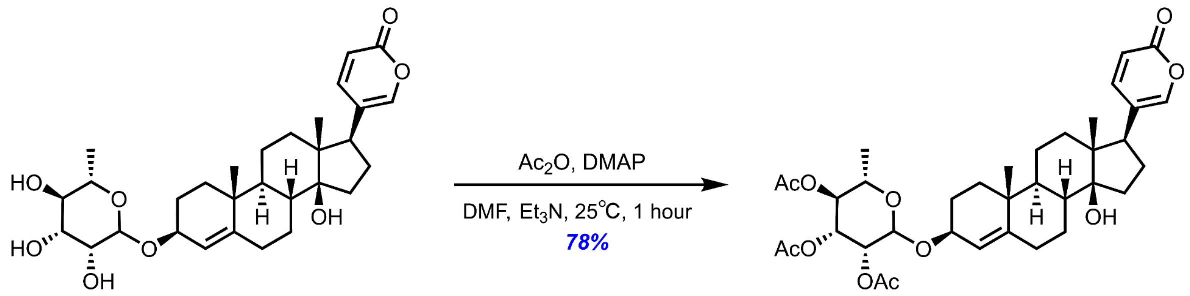


**Chemicals:**
Proscillaridin A (Molekula, 97.5%): used without further purification
4-Dimethylaminopyridine (DMAP) (Sigma Aldrich, ≥99%): used without further purification
Acetic Anhydride (Chemsavers, 97%): used without further purification
N,N-dimethylformamide (DMF) (Sigma Aldrich, ≥99%): used without further purification
Triethylamine (Sigma Aldrich, ≥99.5%): used without further purification

**Procedure:**
To an oven-dried 25 mL oven-dried round bottom flask equipped with a Teflon magnetic stir bar was added proscillaridin A (500 mg, 0.94 mmol, 1 eq) and 4-dimethylaminopyridine (DMAP) (104 mg, 0.85 mmol). To this, N,N-dimethylformamide (7.0 mL) and triethylamine (2.0 mL) were added via syringe, and acetic anhydride (2.0 mL) was added last drop-wise. The reaction was kept at room temperature and stirred for 1 hour, at which point it was determined to be complete by TLC analysis. The reaction mixture was concentrated *in vacuo,* and the crude residue was purified via flash chromatography (0% → 50% EtOAc/Hex, 4.0 cm dia. column), to afford the title compound **4** as a yellow powder (482 mg, 77.9% isolated yield).

**Characterization Data of proscillaridin triacetate**

**TLC:** $R_f$ = 0.50 (70% EtOAc / 30% Hexane), UV active, blue spot by p-anisaldehyde

**ESI-MS:** calculated for $[C_{36}H_{50}O_{11}]^+$ $[M+H]^+$: 658.335; found: 658.342

**FT-IR (ATR, cm$^{-1}$):** 3502, 2938, 1744, 1720, 1662, 1635, 1540, 1445, 1370, 1244, 1222, 1122, 1076, 1046, 1020, 981, 950, 904, 834, 793, 735, 632, 621, 603, 576, 566, 560

**$^1$H NMR of proscillaridin triacetate:** (400 MHz, $CDCl_3$) δ 7.84 (dd, J = 9.8, 2.6 Hz, 1H), 7.21 (dd, J = 2.4, 1.2 Hz, 1H), 6.24 (dt, J = 9.7, 1.3 Hz, 1H), 5.32 – 5.29 (m, 1H), 5.29 – 5.27 (m, 1H), 5.19 (dt, J = 3.2, 1.5 Hz, 1H), 5.04 (td, J = 9.9, 1.4 Hz, 1H), 4.88 (t, J = 1.5 Hz, 1H), 4.21 – 4.03 (m, 1H), 4.03 – 3.92 (m, 1H), 3.77 – 3.64 (m, 1H), 3.51 (ddt, J = 5.8, 4.6, 1.1 Hz, 1H), 3.46 (tt, J = 6.7, 1.2 Hz, 1H), 2.52 – 2.38 (m, 1H), 2.13 (s, 3H), 2.11 – 2.04 (m, 2H), 2.02 (s, 3H), 1.96 (s, 3H), 1.95 – 1.86 (m, 2H), 1.77 – 1.69 (m, 2H), 1.63 (d, J = 8.6 Hz, 1H), 1.61 – 1.56 (m, 2H), 1.56 – 1.52 (m, 1H), 1.52 – 1.42 (m, 2H), 1.35 (dddt, J = 8.5, 7.3, 6.0, 1.3 Hz, 2H), 1.32 – 1.28 (m, 1H), 1.26 (td, J = 2.5, 1.2 Hz, 1H), 1.18 (dt, J = 6.4, 1.1 Hz, 3H), 1.03 (s, 3H), 0.71 (s, 3H).

**$^{13}$C NMR of proscillaridin triacetate:** (101 MHz, $CDCl_3$) δ 170.27, 170.04, 170.02, 162.44, 148.53, 147.26, 146.93, 122.76, 120.34, 115.24, 96.75, 84.95, 84.94, 74.85, 71.72, 71.33, 71.06, 70.49, 69.16, 66.42, 61.79, 60.38, 51.04, 50.09, 48.23, 42.59, 40.62, 37.39, 35.33, 32.63, 32.18, 31.68, 28.69, 28.55, 26.86, 21.22, 21.02, 20.96, 20.79, 20.73, 19.25, 18.89, 17.36, 16.51, 14.17, 13.89.

**$^1$H NMR of proscillaridin triacetate:** (400 MHz, $CDCl_3$)

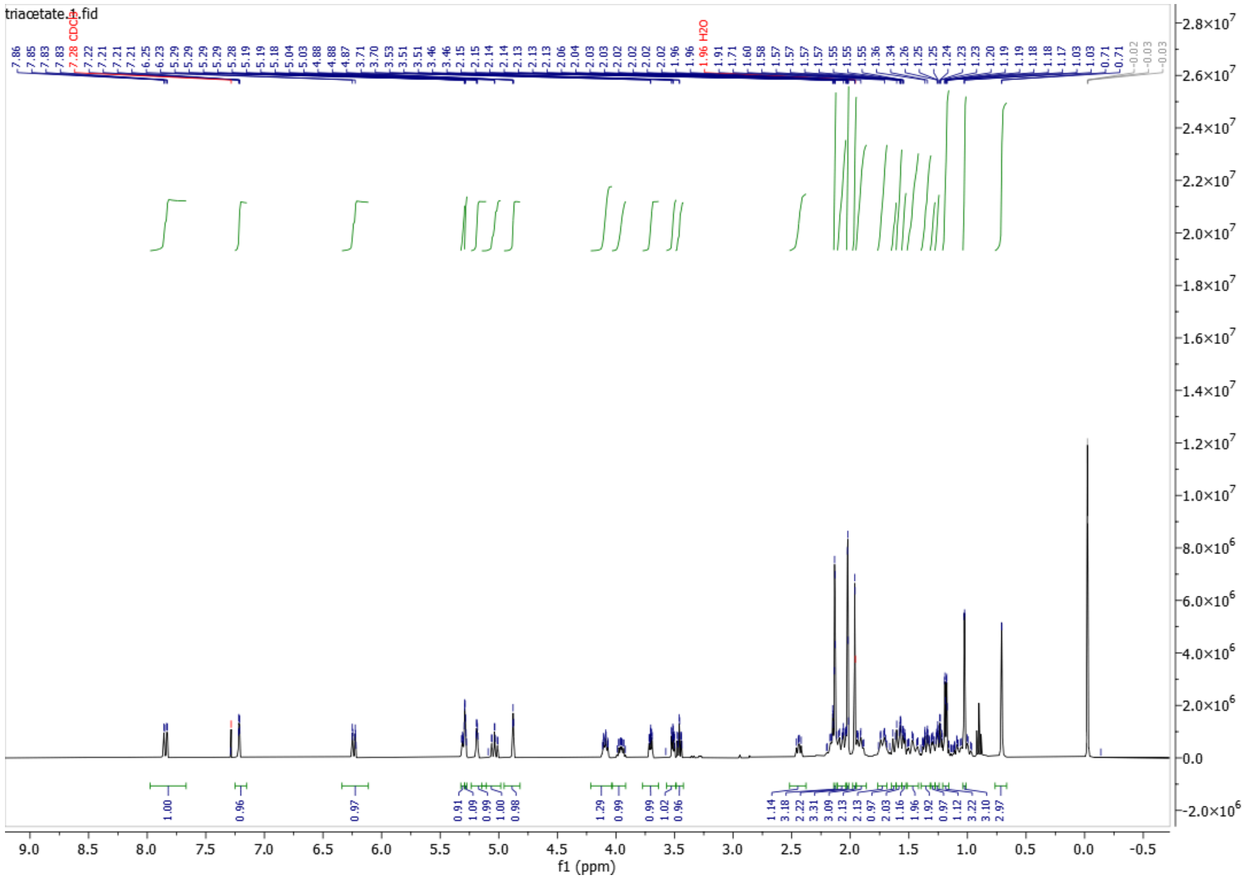

**$^{13}$C NMR of proscillaridin triacetate:** (101 MHz, $CDCl_3$)

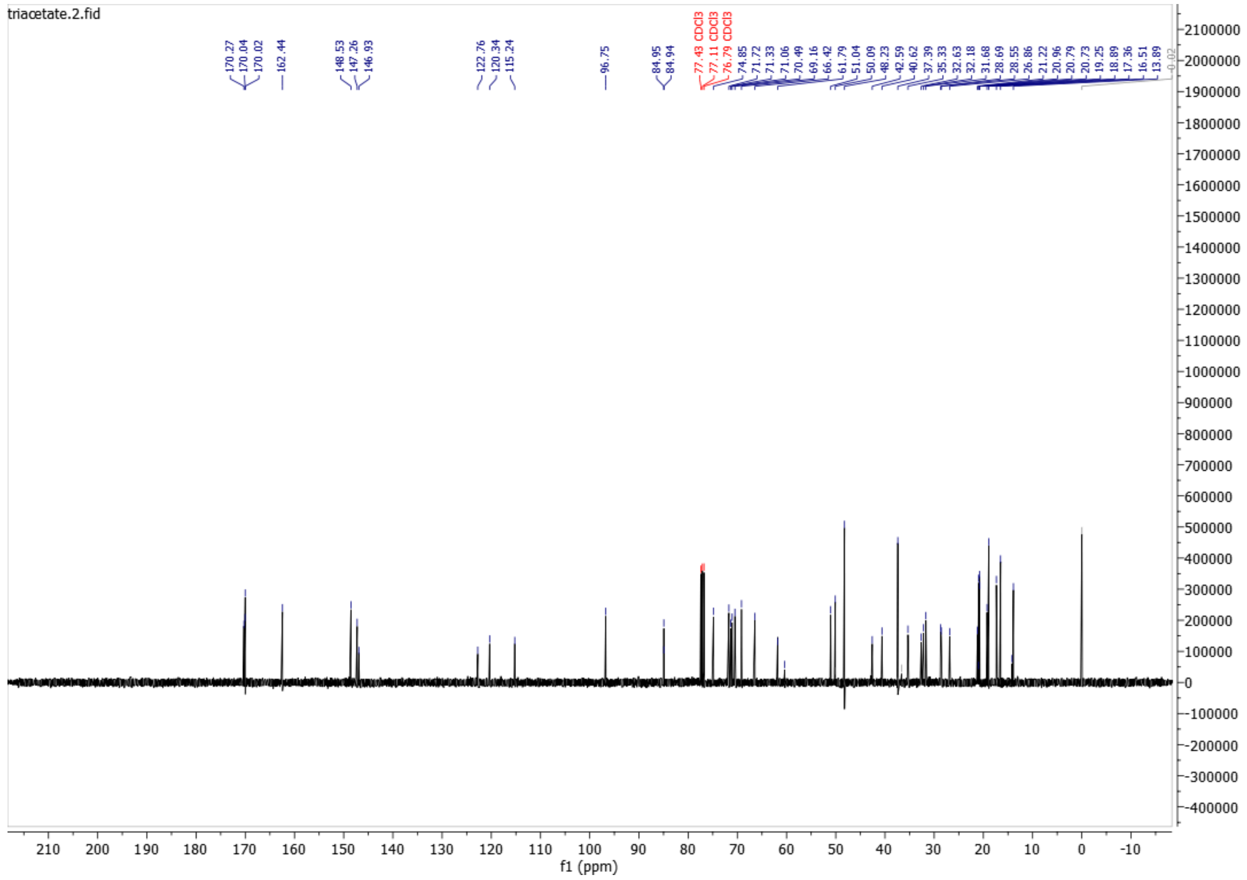

**Preparation of proscillaridin acetonide**

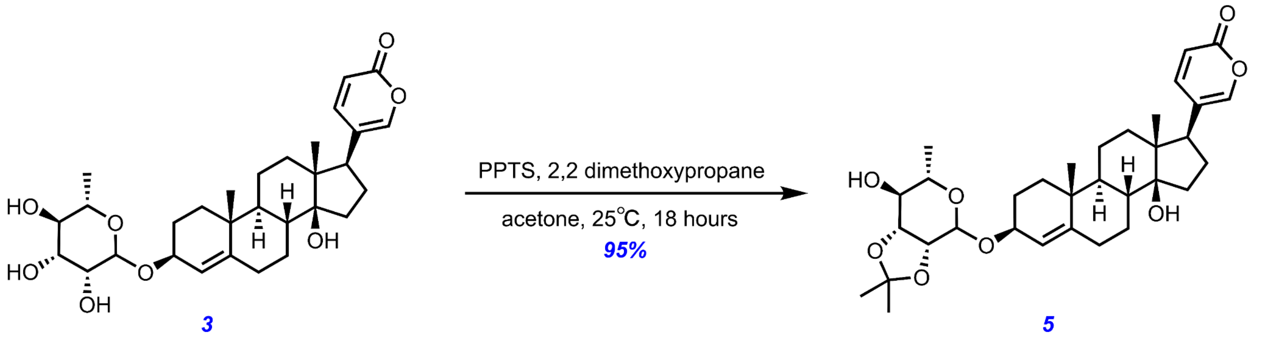


**Chemicals:**
Proscillaridin A (Molekula, 97.5%): used without further purification
2,2 dimethoxypropane (AK Scientific, 98%): used without further purification
Pyridinium p-toluenesulfonate (PPTS) (AK Scientific, 98%): used without further purification
Acetone (Mallinckrodt, 99.9%): used without further purification

**Procedure:**
To an oven-dried 25 mL oven-dried round bottom flask equipped with a Teflon magnetic stir bar was added proscillaridin A (500 mg, 0.94 mmol, 1 eq), 2,2 dimethoxypropane (0.58 mL, 4.70 mmol, 5 eq) via syringe, pyridinium p-toluenesulfonate (PPTS) (24 mg, 0.094 mmol, 0.1 eq) and acetone (16 mL). The reaction was kept at room temperature and stirred for 18 hours, at which point it was determined to be complete by TLC analysis. The reaction mixture was concentrated *in vacuo,* and the crude residue was purified via flash chromatography (0% → 50% EtOAc/Hex, 4.0 cm dia. column), to afford the title compound **5** as a white powder (508 mg, 95.0% isolated yield).

**Characterization Data of proscillaridin acetonide**

**TLC:** $R_f$ = 0.50 (70% EtOAc / 30% Hexane), UV active, blue spot by p-anisaldehyde

**ESI-MS:** calculated for $[C_{33}H_{46}O_8Na]^+$ $[M+Na]^+$: 593.309; found: 593.314

**FT-IR (ATR, $cm^{-1}$):** 3447, 2934, 1708, 1634, 1539, 1448, 1371, 1243, 1219, 1170, 1128, 1071, 1049, 1022, 993, 951, 859, 834, 788, 735, 703, 669, 640, 575, 556

**$^1$H NMR of proscillaridin acetonide:** (400 MHz, DMSO-*d6*) δ 7.93 (dd, J = 9.8, 2.5 Hz, 1H), 7.55 – 7.49 (m, 1H), 6.30 (d, J = 9.7 Hz, 1H), 5.33 (s, 1H), 5.19 (d, J = 6.3 Hz, 1H), 5.07 (s, 1H), 4.27 (s, 1H), 4.10 (dd, J = 9.6, 5.9 Hz, 1H), 4.01 (d, J = 5.7 Hz, 1H), 3.85 (dd, J = 7.4, 5.7 Hz, 1H), 3.57 – 3.45 (m, 1H), 3.06 (dt, J = 9.8, 6.9 Hz, 1H), 2.46 (dd, J = 9.6, 6.4 Hz, 1H), 2.06 (tq, J = 17.5, 8.0, 5.5 Hz, 4H), 1.92 – 1.78 (m, 2H), 1.74 – 1.67 (m, 1H), 1.64 – 1.51 (m, 3H), 1.51 – 1.44 (m, 1H), 1.41 (s, 3H), 1.35 (dd, J = 12.4, 3.4 Hz, 1H), 1.30 (t, J = 3.7 Hz, 1H), 1.27 (s, 3H), 1.26 – 1.16 (m, 2H), 1.12 (d, J = 6.3 Hz, 3H), 0.97 (s, 4H), 0.64 (s, 3H).

**$^{13}$C NMR of proscillaridin acetonide:** (101 MHz, DMSO-*d6*) δ 161.82, 161.81, 149.71, 147.82, 146.94, 146.92, 123.12, 121.09, 114.71, 108.49, 96.03, 83.59, 78.56, 76.27, 74.01, 73.35, 66.20, 50.42, 50.04, 48.38, 41.95, 37.41, 35.45, 32.44, 32.30, 28.89, 28.49, 27.26, 26.73, 21.36, 19.03, 17.77, 17.08.

**$^1$H NMR of proscillaridin acetonide:** (400 MHz, DMSO-*d6*)

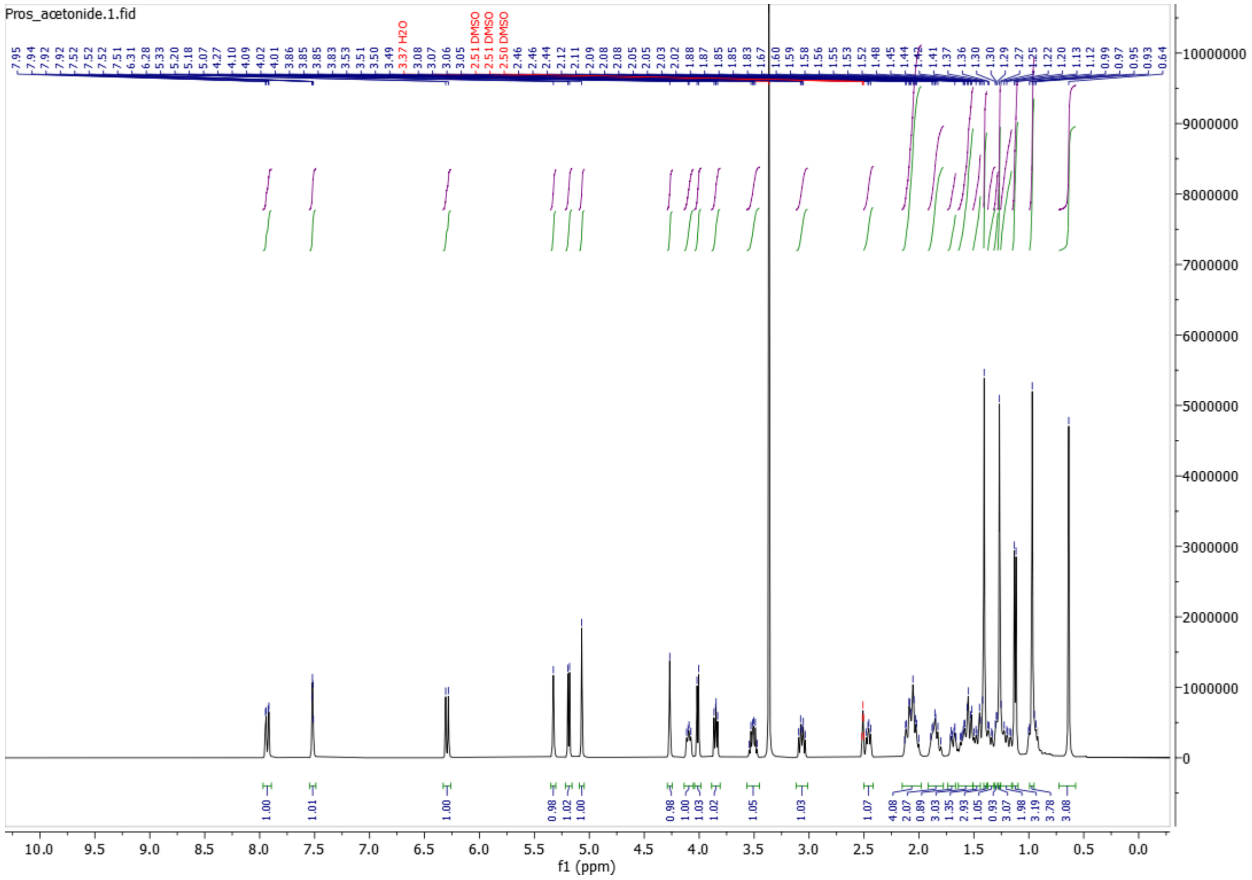

**$^{13}$C NMR of proscillaridin acetonide:** (101 MHz, DMSO-*d6*)

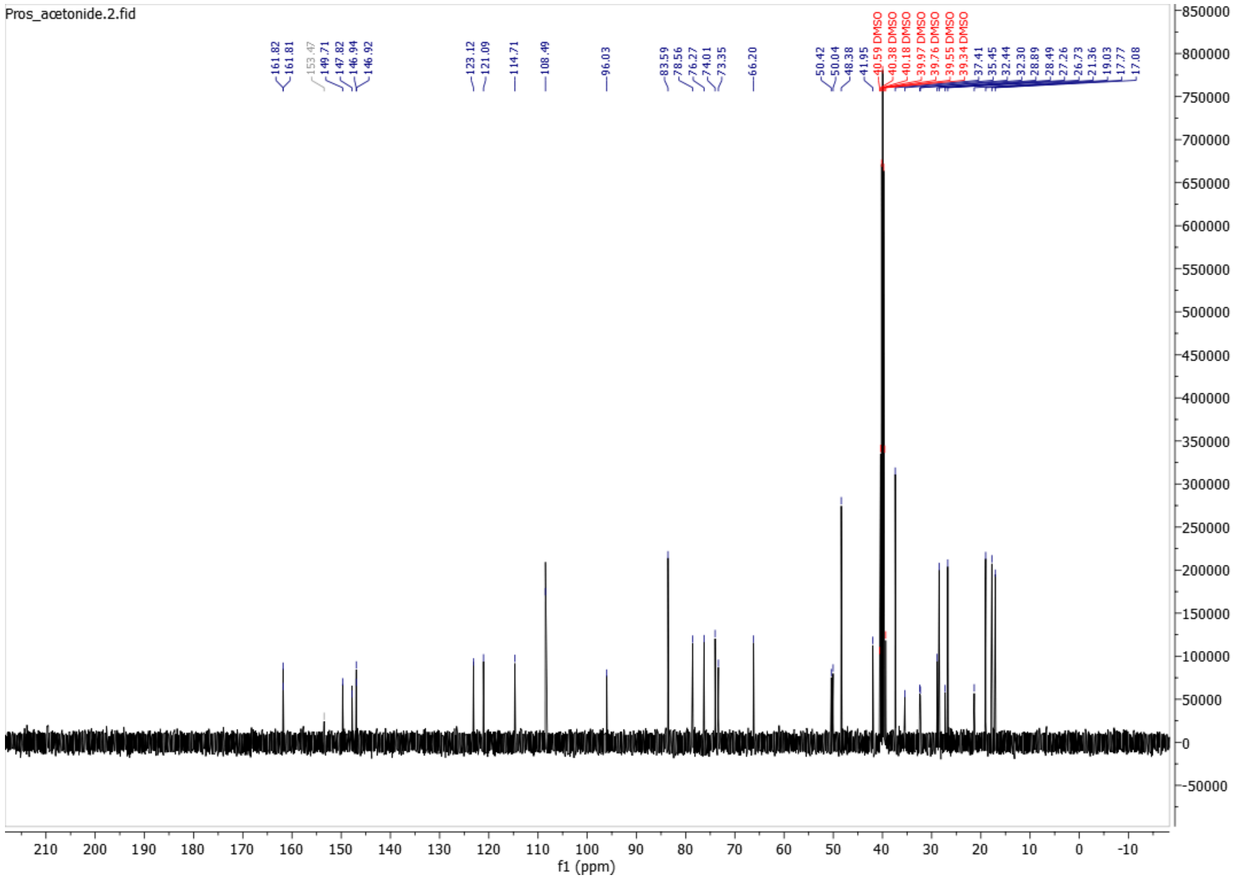

#### b. Cell Culture

HCT-116 and HT-29 human colorectal cancer cell lines were obtained from the European Collection of Authenticated Cell Cultures (ECACC) and cultured in McCoy's 5A Medium (Tribioscience) supplemented with 10% v/v fetal bovine serum (FBS, Gibco) and 1% v/v 100x penicillin-streptomycin (Tribioscience). CT26, a murine colorectal carcinoma cell line, was obtained from American Type Culture Collection (ATCC), and cultured in RPMI-1640 Medium (Tribioscience) supplemented with 10% v/v fetal bovine serum (FBS, Gibco) and 1% v/v 100x penicillin-streptomycin (Tribioscience). HepG2, a human liver cancer cell line, was obtained from the European Collection of Authenticated Cell Cultures (ECACC) and cultured in High Glucose Dulbecco's Modified Eagle Medium (DMEM) (Hygia Reagents) supplemented with 10% v/v fetal bovine serum (FBS, Gibco) and 1% v/v 100x penicillin-streptomycin (Tribioscience). A human embryonic kidney cell line, HEK-Blue™ TNF-α, stably transfected with a TNF-α inducible secreted alkaline phosphatase (SEAP) reporter gene, was obtained from Invivogen (Cat. #hkb-tnfdmyd) and cultured in High Glucose Dulbecco's Modified Eagle Medium (DMEM) (Hygia Reagents) supplemented with 10% v/v fetal bovine serum (FBS, Gibco) and 1% v/v 100x penicillin-streptomycin (Tribioscience). HEK-BlueTM CD40L cells engineered to express the CD40 receptor were obtained from Invivogen (Cat. #hkb-cd40), and cultured in High Glucose Dulbecco's Modified Eagle Medium (DMEM) (Hygia Reagents), supplemented with 10% v/v fetal bovine serum (FBS, Gibco) and 1% v/v 100x penicillin-streptomycin (Tribioscience). HEK-Blue™ IFN-γ cells engineered to monitor human type II interferon IFN-γ-induced STAT1 stimulation were obtained from Invivogen (Cat. #hkb-ifng) and cultured in High Glucose Dulbecco's Modified Eagle Medium (DMEM) (Hygia Reagents), supplemented with 10% v/v fetal bovine serum (FBS, Gibco) and 1% v/v 100x penicillin-streptomycin (Tribioscience). HEK-Blue™ IL-10 cells engineered for the detection of human IL-10 to monitor JAK1/STAT3 signaling were obtained from Invivogen (Cat. #hkb-il10), and cultured in High Glucose Dulbecco's Modified Eagle Medium (DMEM) (Hygia Reagents), supplemented with 10% v/v fetal bovine serum (FBS, Gibco) and 1% v/v 100x penicillin-streptomycin (Tribioscience). HEK-Blue™ IL-2 cells engineered for the detection of human IL-2 to monitor JAK1/3-STAT5 signaling were obtained from Invivogen (Cat. #hkb-il2), and cultured in High Glucose Dulbecco's Modified Eagle Medium (DMEM) (Hygia Reagents), supplemented with 10% v/v fetal bovine serum (FBS, Gibco) and 1% v/v 100x penicillin-streptomycin (Tribioscience). LEADING LIGHT® Wnt Reporter 3T3 mouse embryonic fibroblast cells were obtained from Enzo Life Sciences (Cat. #ENZ-61002-0001) and cultured in High Glucose Dulbecco's Modified Eagle Medium (DMEM) (Hygia Reagents), supplemented with 10% v/v fetal bovine serum (FBS, Gibco), and 1% v/v 100x penicillin-streptomycin (Tribioscience).

#### d. Flow Cytometry

##### i. *Cell Cycle Flow Cytometry*

HCT-116, HT-29, and CT26 cells were seeded at 70% confluency in 6-well tissue-culture treated plates (Corning) in 1.50 mL of cell-line respective media (McCoy's 5A Medium for HCT-116 and HT-29, RPMI-1640 for CT26) supplemented with 10% v/v fetal bovine serum (FBS, Gibco) and 1% v/v 100x penicillin-streptomycin (Tribioscience). Cells were incubated for 24 hours at 37ºC (5.0% $CO_2$) to allow for attachment, after which 7.5 µL of 200-fold dilutions of compounds **1** through **5** were added to achieve a final concentration of 100 nM. A negative control of 0.5% v/v DMSO was also included. After 24 hours of incubation with the drug, the supernatant media was collected in 2 mL Eppendorf tubes (Eppendorf) and the cells were trypsinized with 400 µL of 0.25% EDTA-trypsin (Tribioscience) per well. The cells were incubated for 10 minutes, and the trypsin was deactivated with 300 µL of cell-line respective media. The detached cells were moved into the same Eppendorf tubes as those containing their drugged media and centrifuged for 9 minutes at 2500 rpm. The supernatant was removed, and the cell pellet was washed in 500 µL of 1x PBS and re-centrifuged for 10 minutes at 3500 rpm. The PBS was aspirated off and the cells were resuspended in 500 µL of 70% ethanol at 4ºC for 30 minutes, then re-pelleted by centrifugation. The ethanol was aspirated off, the cells were resuspended in 25 µL of ribonuclease A (1 mg/mL in DI water, Research Products International, 97% purity) and 100 µL of PBS and were then incubated for 5 minutes at 37ºC (5.0% $CO_2$). The cells were then stained in the dark with 1 µL propidium iodide (1 mg/mL in DI water, AK Scientific, 98%) for 30 minutes. DNA content was then analyzed to determine cell cycle stage with a BD Accuri C6 Flow Cytometer. Each sample was run to 50,000 events, and single cells were gated to exclude debris based on forward scatter and side scatter, followed by FL2-A channel gating. Results were analyzed using FlowJo Software 10 to determine the percentage of cells in each stage of the cell cycle ($G_0$/$G_1$, S, $G_2$/M).

MFI was calculated against FL1-A and expressed as percentages in relation to the control. Results were analyzed using FlowJo Software 10.

#### e. Reporter Assays

**i.** ***NF-κB (TNF-α) Reporter Secreted Alkaline Phosphatase Cell Assay***

### 3. Supplementary Figures

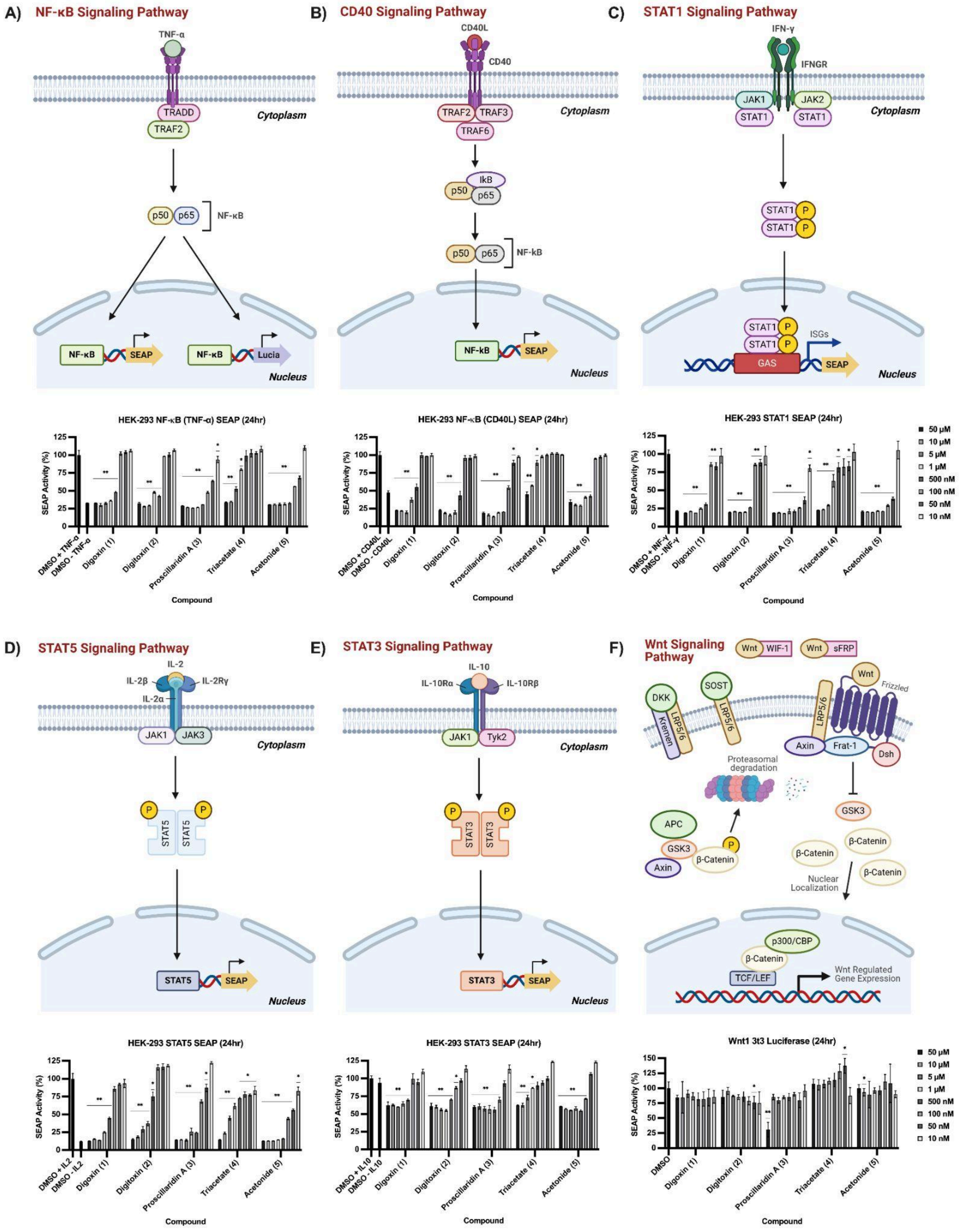

**S1: Reporter cell signal pathways and profiling of activity of compounds through secreted alkaline phosphatase (SEAP) and Wnt1 luciferase cell assays.** A negative control was established using 0.5% v/v DMSO. Data is represented as means (relative to a background control for Wnt1 luciferase) ± SD compared with the negative control using a Welch's t-test ($n = 4$) (*$P < 0.05$, **$P < 0.01$).
